# Regulation of the human voltage-gated proton channel by membrane sterols

**DOI:** 10.64898/2026.08.20.746042

**Authors:** Shuo Han, Rui Duan, Sarah Applewhite, Shuyi Wang, Grace Wang, Mingxing Qian, Douglas F. Covey, Xiaoqin Zou, Shizhen Wang

**Affiliations:** Division of Biological and Biomedical Systems School of Science and Engineering University of Missouri-Kansas City, Kansas City, MO, USA; Department of Physics and Astronomy, University of Missouri, Columbia, MO, USA; Department of Biochemistry, University of Missouri, Columbia, MO, USA; Dalton Cardiovascular Research Center, University of Missouri, Columbia, MO, USA; Institute for Data Science and Informatics, University of Missouri, Columbia, MO, USA; Department of Developmental Biology, Washington School of Medicine, St. Louis, MO, USA; Taylor Family Institute for Innovative Psychiatric Research, Washington University School of Medicine, St. Louis, MO, USA

## Abstract

Cholesterol is a key component of eukaryotic cell membranes, promoting membrane stability and modulating the function of many membrane proteins, including ion channels. In our previous work using purified human voltage-gated proton channel proteins, we showed that cholesterol inhibits the hHv1 channel by altering the conformational dynamics of its S4 segment, the key element that senses membrane voltage to control proton permeation. In the present work, we examined the effects of cholesterol analogs and potential sites in the hHv1 channel mediating cholesterol inhibition using site-directed mutagenesis and docking simulations. Our results showed that desmosterol, the immediate precursor of cholesterol, markedly attenuates cholesterol inhibition. Using single-molecule Fluorescence Resonance Energy Transfer (smFRET), we showed that desmosterol attenuates cholesterol inhibition by promoting the intermediate and open state conformations of the S4 segment. Moreover, we identified multiple residues in the hHv1 channel that are critical for cholesterol inhibition, including Y141A in the S2 segment, which reduces cholesterol inhibition by nearly 3-fold. Our smFRET results showed that the Y141A mutation promotes the intermediate conformation in the S4 segment, which underlies the attenuation of cholesterol inhibition. Consistently, docking simulations also revealed multiple residues spanning the transmembrane domain, rather than clustered within a single localized pocket. Our work identified the key molecular determinant in the hHv1 channel that mediates cholesterol inhibition and also provided a mechanism linking the conversion between demosterol and cholesterol by DHCR24 to pH homeostasis in many cells, such as phagocytes, cardiomyocytes, neurons and microglial cells.

## INTRODUCTION

Voltage-gated proton (Hv) channels are standalone voltage sensors without the pore-forming domain commonly seen in canonical voltage-gated cation channels (1, 2). The voltage dependence of Hv channels is remarkably shifted upon both intra- and extracellular pH, normally mediating proton efflux (3). In addition to voltage and pH, Hv channels are also modulated by temperature (4), mechanical forces (5, 6) and small molecules like Zn^2+^ (7, 8), polyunsaturated fatty acid (9), albumin (10), ATP (11) and even an auxiliary subunit (12). Like other canonical voltage sensors, the S4 transmembrane segment of Hv channels carries multiple positively charged residues, serving as the primary element sensing electrical voltage (13, 14). Moreover, the S4 segment is also involved in the regulation of Hv channels by Zn^2+^, temperature and mechanical forces (4, 5, 8). Recently, using purified human Hv1 (hHv1) proteins reconstituted into liposomes with defined lipid compositions, we showed that membrane cholesterol inhibits the hHv1 channel by stabilizing its S4 segment at resting state conformations (15). However, regulation of ion channels by membrane cholesterol could be through the direct, specific interaction or by altering the physical properties of membranes (16). A most recent study suggested that activation of hHv1 channels by depleting membrane cholesterol might be associated with disruption of cholesterol-rich membrane domains (17). In phagocytes, the hHv1 channels work with NADPH oxidase to facilitate the production of reactive oxygen species to destroy invasive pathogens, a critical process that the innate immunity relies on (18, 19). Interestingly, the NADPH oxidase complex is dependent on membrane cholesterol for assembly and function (20), and hHv1 channels were also identified in cholesterol-rich lipid raft microdomains in human T lymphocytes and sperm (21, 22). So, defining the mechanisms underlying cholesterol regulation of hHv1 channels will be critical to understand their function in human phagocytes, sperm and many other cells.

In the present work, we examined the functional effects of cholesterol analogs on the hHv1 channel and also performed mutagenesis and docking simulations to explore the mechanisms underlying cholesterol inhibition. We used the cholesterol enantiomer, ent-cholesterol, to distinguish specific cholesterol-protein interactions from nonspecific effects of cholesterol, since ent-cholesterol has the same effects as natural cholesterol on the physical properties of membranes (23). Therefore, if natural and ent-cholesterol have different effects on the function of hHv1 channels, this may suggest inhibition via direct binding (24). Our liposome flux assay data showed that ent-cholesterol is only slightly less potent than natural cholesterol in inhibiting hHv1 channels. Excitingly, we found that the cholesterol hHv1 interactions are very specific: desmosterol, which differs from cholesterol only by a double bond in the side chain, reverses cholesterol inhibition. Using site-directed mutagenesis and docking simulations, we identified multiple residues involved in cholesterol inhibition spanning the entire transmembrane domain. We further showed that the alanine substitution at a key residue, Y141, shifts the half-inhibition concentration by nearly 3-fold. Collectively, our results suggested that cholesterol may inhibit hHv1 channels through direct binding at multiple sites, and that cholesterol metabolism may play a role in regulating the function of hHv1 channels in many cells.

## MATERIALS AND METHODS

### Protein mutation expression, purification, and fluorophore labeling

Human voltage-gated proton channels (hHv1) carrying an N-terminal 6xHis-tag were expressed in *E. coli* host cells and purified by immobilized metal affinity chromatography as described previously (13, 15, 25). The cysteine residues introduced at either K125C-S224C or K169C-Q194C of hHv1 channel proteins, one pair at a time, and cysteine mutants for smFRET studies carry the additional N214R background mutation that prevents proton accumulation into liposomes (13, 15, 26). The purified cysteine mutant proteins were conjugated to modified Cy3 and Cy5 fluorophores with improved photostability carrying the maleimide reactive group (27) and then reconstituted into liposomes immediately for smFRET imaging.

### Liposome reconstitution and liposome flux assay

For the liposome flux assay, the purified hHv1 WT or mutant proteins were reconstituted into POPE/POPG (3/1, w/w) liposomes at a protein-lipid ratio of 1:200 (w/w). The hHv1 liposomes were used immediately or stored at −80 °C for later use. The K^+^ gradient across liposomes was established as described previously (15). The quenching of ACMA fluorescence by proton uptake into liposomes through either hHv1 channels or proton ionophore CCCP was monitored with a 96-well fluorescence plate reader, with an excitation wavelength of 390 nm and emission wavelength of 460 nm. After reading the initial ACMA fluorescence for 4 min, the K+ ionophore valinomycin at a final concentration of 0.45 µM was added to generate intraliposomal negative electrical potentials to drive proton uptake, and ACMA fluorescence readings were resumed with the same optical settings for ∼15 min. At the end of the liposome flux assay, proton ionophore CCCP was added as a positive control. The hHv1 WT liposomes were included in every batch of liposome flux assays for normalization (13). The relative activities of hHv1 channels were calculated using the following equation:

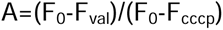

Where F_0_, F_val_, and F_cccp_ were the ACMA fluorescence at initial, after adding valinomycin or CCCP, respectively.

### smFRET imaging and data analysis

For smFRET imaging, the fluorophore-labeled hHv1 proteins were reconstituted into POPE/POPG (3/1, w/w) liposomes at a very low protein-lipid ratio of 1:4,000 (w/w) so that most liposomes were either empty or contained only one hHv1 channel protein (13, 15). Slides and coverslips were coated with PEG/biotin-PEG and then assembled into flow chambers as described by Joo et al. (28). The biotinylated anti-His-tag antibodies were attached to biotin-PEG coverslip surfaces through neutravidin, and hHv1 liposomes with cytosolic 6*His tag facing outside were selectively retained on the coverslip surface for smFRET imaging. All smFRET data were performed under −85 mV liposome potential, generated by transliposomal K^+^ gradient (5 mM inside and 150 mM outside), in the presence of 0.45 µM valinomycin (13, 15). All smFRET imaging movies were collected with a customized objective-based TIRF microscope or a Nanoimager of Oxford Nanoimaging using a 532 nm laser line (∼1.0 W/cm^2^). The smFRET traces were extracted from movies using the SPARTAN software package developed by the Blanchard group (29), and then further manually picked to ensure the data quality, with previously established criteria, including FRET Lifetime >50 frames, minimum mean FRET >0.15, donor/acceptor correlation coefficient between −1.1 and 0.5, signal-to-noise ratio >8, background ground noise <70, donor blinks <4, and overlap molecule-removal function checked (13).

All smFRET traces were idealized with a kinetic model containing low, medium and high FRET states and an additional zero FRET state accounting for FRET events after photobleaching using the Maximum Point Likelihood (MPL) algorithm (30). FRET histograms and contour maps were generated from all idealized smFRET traces in each group using a bin size of 0.03. FRET state occupancies were calculated as the fractional areas under the corresponding peaks in the histograms. The uncertainties were estimated by bootstrapping the idealized smFRET traces 1,000 times and determining the fractional occupancy of each FRET state for each resampled dataset. Equilibrium constants between different FRET states were calculated from ratios of state populations and presented as lnK values, along with error bars reflecting propagated uncertainties in the state-occupancy data.

### Computational approaches for prediction of cholesterol-hHv1 complexes

Because no experimentally determined structure of a cholesterol-bound hHv1 complex is currently available, AlphaFold3 (version 3.0.1) (31) was employed to predict potential cholesterol-binding modes. The amino acid sequence of hHv1 and the SMILES representation of cholesterol were provided as inputs. Structure prediction was performed using the default AlphaFold3 settings with 20 random seeds, generating a total of 100 cholesterol-hHv1 complex models. To reduce structural redundancy, the predicted complexes were clustered based on ligand root mean square deviation (L-RMSD) using a cutoff of 5 Å. For each cluster representative, protein residues with at least one heavy atom within 6.0 Å of cholesterol were identified. Residues frequently observed in contact with cholesterol across the predicted models were selected for further analysis.

To complement the AlphaFold3 predictions, molecular docking was performed using representative open and closed hHv1 conformations obtained from molecular dynamics (MD) simulations reported in a previous study (32). Docking calculations were carried out using Glide (33), implemented in the Schrödinger Suite (release 2025-3) (34). The receptor grid was defined to encompass the entire transmembrane domain of hHv1. Cholesterol was docked using the standard precision (SP) protocol with default parameters. Protein residues within the predicted cholesterol-binding regions were identified from the top-ranked docking poses and combined with the AlphaFold3-predicted residues for subsequent experimental validation by site-directed mutagenesis.

## RESULTS

### Membrane sterols have differential effects on the function of hHv1 channels

By reconstituting purified proteins into liposomes with defined lipid compositions, our previous studies showed that cholesterol strongly inhibits hHv1 channels(15). However, the mechanism of cholesterol inhibition is yet to be elucidated. In the present work, we further examined the effects of desmosterol, the immediate precursor to cholesterol, and 2 cholesterol stereoisomers (Fig. 1a), epi-cholesterol and ent-cholesterol. Among them, ent-cholesterol is an enantiomer of cholesterol, which has the same effects on membrane physical properties as cholesterol (35–37). By reconstituting hHv1 channels into POPE/POPG (3/1, w/w) liposomes containing 30% of cholesterol, epi-cholesterol and ent-cholesterol, our liposome flux assay results showed that hHv1 channels were completely inhibited by both 30% cholesterol and epi-cholesterol (Fig. 1b, c). We found that the hHv1 channel exhibits very low enantioselectivity, showing only residual channel activity in the presence of 30% ent-cholesterol (Fig. 1b, c). However, 30% of desmosterol does not have any significant inhibitory effects on hHv1 channels, although it only contains an additional double bond in the side chain between carbon atoms 24 and 25, suggesting a potential specific interaction between cholesterol and hHv1 channels (Fig. 1a). To our surprise, in the presence of 30% cholesterol (w/w), addition of 10% desmosterol almost completely abolishes cholesterol inhibition, suggesting that desmosterol and cholesterol may either have shared binding sites or have differential structural effects on hHv1 channels (Fig. 1d).

**Figure 1:**
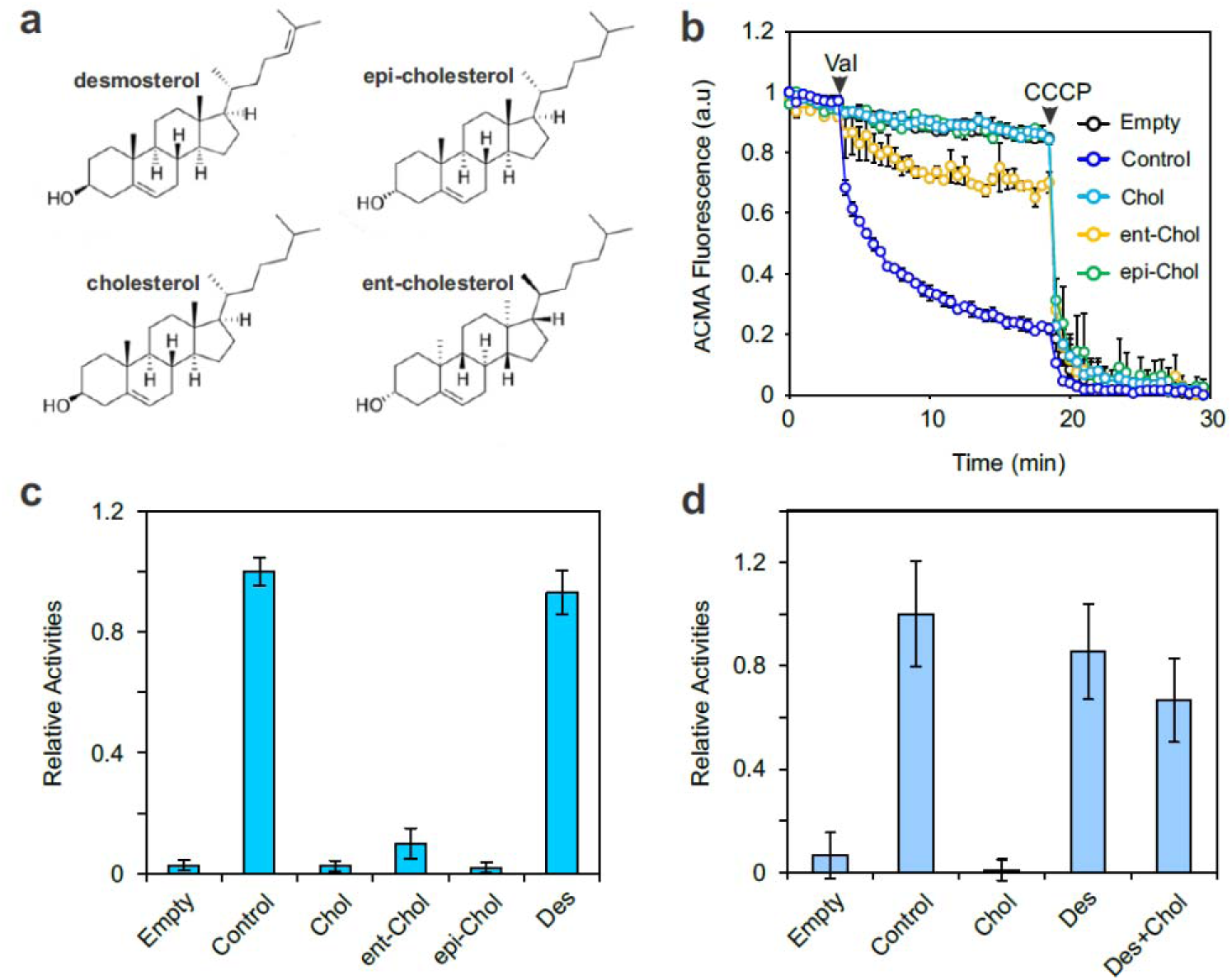
Membrane sterols have different effects on the function of hHv1 channels. **a**. Chemical structures of desmosterol, cholesterol, epi-cholesterol (diastereomer) and ent-cholesterol (enantiomer) (37). **b**. Liposome flux assay curves of hHv1 channels reconstituted into liposomes containing 30% cholesterol, ent-cholesterol and epi-cholesterol. Liposomes without hHv1 channels (empty) or with hHv1 channels without cholesterol (control) were included. All data were presented as mean±SE, n=3. **c**. The relative activities of hHv1 channels reconstituted into liposomes containing 30% (w/w) of different membrane sterols. The enantiomer of cholesterol (ent-Chol) is slightly less potent than cholesterol (Chol) in inhibiting hHv1 channels, while its immediate precursor molecule, desmosterol (Des), does not demonstrate significant inhibitory effects. All data were presented as mean±SE, n=3-6. **d**. The relative activities of hHv1 channels reconstituted into liposomes with 30% (w/w) cholesterol (Chol), 10% (w/w) desmosterol (Des) or 30% cholesterol and 10% desmosterol (Des+Chol). The hHv1 channel was almost completely inhibited by 30% cholesterol, a response that was almost completely reversed by 10% desmosterol. All data were presented as mean±SE, n=3-6.

### Desmosterol promotes open conformations of the hHv1 S4 segment

Our previous work already established that cholesterol inhibits hHv1 channels by stabilizing the S4 segment in its resting state (15). To understand how desmosterol reverses cholesterol inhibition, we examined its effects on the structural dynamics of the S4 segment using smFRET. Our previous work showed that the S4 movements can be monitored by measuring FRET changes between Cy3 and Cy5 fluorophores conjugated to either the K125C-S224C or K169C-Q194C labeling sites (Fig. 2a). In the present work, we performed FRET measurements at −85 mV with the rationale that the activating effects of demosterol would be detectable on hHv1 channels in resting membrane potential. As reported previously, the S4 segment exhibited 3 major conformational populations, as shown by smFRET data collected from two different labeling sites. As expected, the effects of 20% cholesterol were minimal, since hHv1 channels were exposed to the resting membrane voltage of −85 mV (Fig. 2b, d). The addition of 10% desmosterol, however, remarkably enriched the high-FRET population and diminished the low-FRET population at the K125C-S224C labeling sites and consistent changes were also observed at the K169C-Q194C labeling sites (Fig. 2b, d). The smFRET data from both labeling sites indicate that desmosterol stabilizes the S4 segment at a more outward conformation, thus promoting channel opening. As shown by contour maps, in hHv1 proteoliposomes with 20% cholesterol, the addition of 10% desmosterol almost eliminated the conformational changes induced by 20% cholesterol, which matchs with its functional effects in reversing cholesterol inhibition. However, we did not observe remarkable activation effects in hHv1 proteoliposomes containing only desmosterol in our liposome flux assays, perhaps because hHv1 channels are already sufficiently active to saturate assays. Since demosterol is the immediate precursor of cholesterol, our findings suggest that cholesterol metabolism may contribute to cellular pH homeostasis by regulating the hHv1 channel.

**Figure 2.**
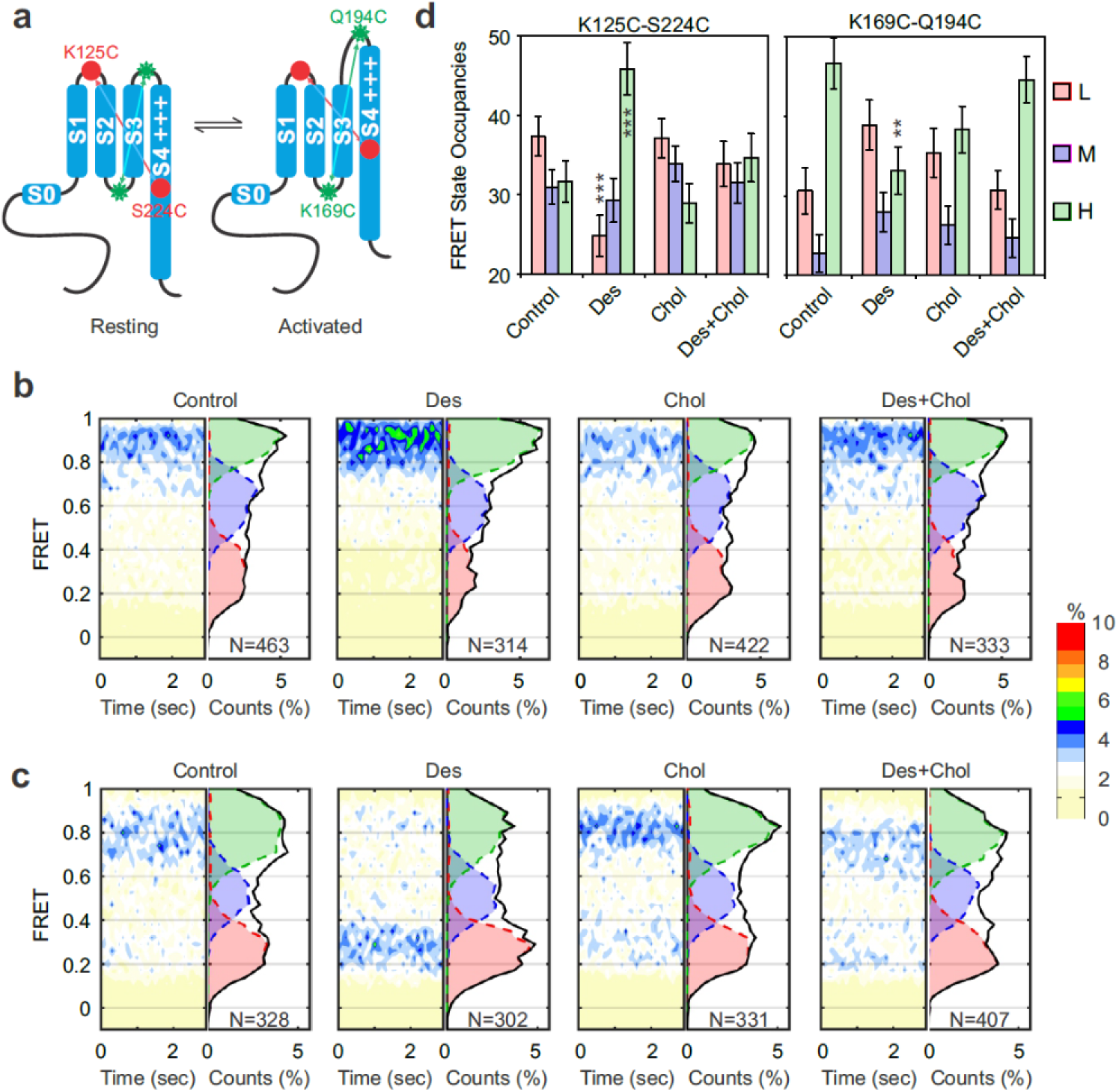
Desmosterol attenuates cholesterol-mediated inhibition of hHv1 channels by reversing the structural effects of cholesterol on the voltage-sensing S4 segment. **a**. Schematic illustration of fluorophore labeling configurations to monitor the conformational transitions of the hHv1 S4 segment using smFRET. Cysteine mutations were introduced at the K125/S224 (red dots) or K169/Q194 (green sun icons) sites, one pair at a time. The outward movements of the S4 segment are reflected by increased FRET between fluorophores conjugated to the K125 and S224 sites, but decreased FRET to the K169 and Q194 sites. **b** and **c**. FRET contour maps and histograms of FRET traces from the K125C/S224C (**b**) and K169C/Q194C (**c**) labeling sites, reconstituted into liposomes without (control) or with 10% desmosterol (Des), 20% cholesterol (Chol) or both (Des+Chol). The N number indicated the total number of FRET traces included in the analysis. **d**. FRET state occupancies of FRET traces from the K125C/S224C and K169C/Q194C labeling sites, in hHv1 proteoliposomes without (control) or with 10% desmosterol (Des), 20% cholesterol (Chol) or both (Des+Chol). A kinetic model containing 3 FRET states, including low (L), medium (M) and high (H) FRET states, was used to idealize FRET traces. The error bars indicate uncertainties in FRET state occupancy data obtained via bootstrapping. Two-tailed Student’s t-tests were performed to examine changes in FRET state occupancies in comparison to the control condition, with * denoting p<0.05, ** denoting p<0.01 and *** denoting p<0.001.

### Mutagenesis and docking simulations suggest multiple cholesterol binding sites

To further elucidate the molecular basis of cholesterol inhibition of hH1 channels, we performed mutagenesis to identify residues in the hHv1 channel essential for cholesterol inhibition. We selected residues similar to cholesterol recognition amino acid (CRAC or CARC) motifs (i.e., L/V-X(_1-5_)-Y-X(_1-5_)-K/R or the inverted sequence) in the hHv1 transmembrane domain. Of the 24 residues subjected to mutational analysis, alanine mutations at 9 residues led to significant decreases in cholesterol inhibition (Fig. 3a). Several mutants, including Y141A, L147A, F149A and L200A, maintain a similar level of proton channel activities, while their inhibition by 20% cholesterol was almost abolished, much more than the Y134A mutant previously identified by electrophysiological studies (17). These 9 residues are located in the S1-S4 segments and span the entire transmembrane domain (Fig. 3b).

**Figure 3.**
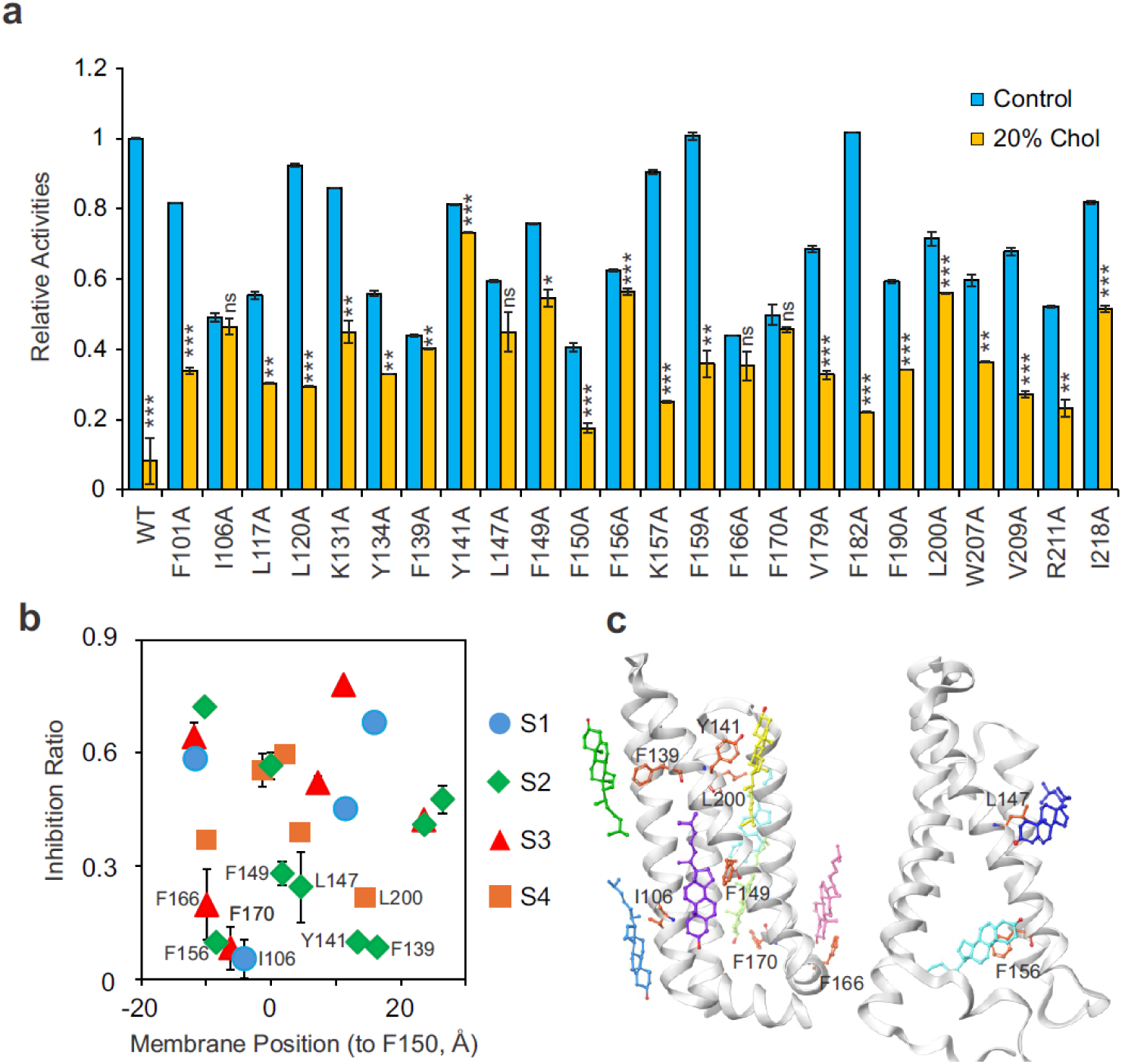
Mutations in the transmembrane domain attenuate cholesterol inhibition of hHv1 channels. **a**. Activities of hHv1 mutants reconstituted into liposomes without (control) or with 20% cholesterol. All data were presented as mean±SE, n=3-9. Two-tailed Student’s t-tests were performed to examine changes in activities of hHv1 WT or mutants without or with 20% cholesterol, with ns denoting p≥0.05, * denoting p<0.05, ** denoting p<0.01 and *** denoting p<0.001. **b**. Ratio of inhibition by 20% cholesterol for hHv1 WT and mutant channels, calculated from data in panel **a**. All data were presented as mean±SE, n=3-9. Residues are differentially displayed by their positions in S1-4 transmembrane segments and their relative positions in membranes with respect to F150 in the S2 segment. Alanine mutations at residues that lead to an inhibition ratio lower than 0.3 were highlighted. **c**. Cholesterol-binding sites identified by computational modeling of hHv1. Left panel, 7 cholesterol-hHv1 complex models generated by AlphaFold3 that identified I106, F139, Y141, F149, F166, F170, and L200 as key residues (shown in orange). All models are aligned to a common hHv1 structure; therefore, the positions of cholesterol and the highlighted residues may differ slightly from those in the individual complex structures. Right panel, a representative cholesterol-binding pose obtained by docking to an MD-derived hHv1 structure, identifying F156 (orange) as a potential cholesterol-interacting residue. For L147, which was experimentally identified as affecting cholesterol inhibition, a representative cholesterol-binding pose was also generated by docking into the MD-derived hHv1 structure.

We also combined AlphaFold3 modeling with molecular docking to identify potential cholesterol-binding sites. Our docking simulations also revealed multiple cholesterol-binding sites with interacting residues distributed throughout the transmembrane domain of the hHv1 channel, rather than clustered within a single localized pocket (Fig. 3c). Among them, alanine mutants of I106, F139, Y141, F149, F156, F170, F166 and L200 residues also demonstrated significant reduction in cholesterol inhibition as revealed by our liposome flux assays (Fig. 3b). Collectively, our mutagenesis and simulation results suggest that cholesterol may inhibit hHv1 through direct binding to multiple sites, rather than a single localized binding pocket.

### Cholesterol inhibition of the hHv1 channel is conformationally dependent

Since the Y141A mutant retains the same level of channel activity as that of WT, while its sensitivity to cholesterol inhibition was almost completely abolished (Fig 3a). We further examined its channel function in liposomes containing 0-50% cholesterol. Dose-dependent inhibition of 141A by the cholesterol mutant was well fit by the Hill equation, yielding a half-inhibition concentration of 31%, corresponding to an approximately threefold shift compared with 11% for the WT (Fig. 4a). We examined the effects of the Y141A mutation on the structural dynamics of the S4 segment using smFRET, using the two labeling sites established in our previous work. All smFRET measurements at a resting voltage of −85 mV to detect potential structural changes induced by the Y141A mutation that may promote channel opening. Again, no significant changes were observed in the WT channel with or without 20% cholesterol (Fig. 4a, b). However, data collected from the mutants carrying the Y141A mutation background showed that the Y141A mutation significantly promoted the low and medium FRET populations at the K169C-Q194C labeling site, and the high and medium FRET populations at the K125C-S224C labeling sites (Fig. 4a, b, c). These results suggested that the Y141A mutation attenuates cholesterol inhibition by stabilizing the S4 segment at pre-activated or activated conformations (Fig. 4d).

**Fig 4.**
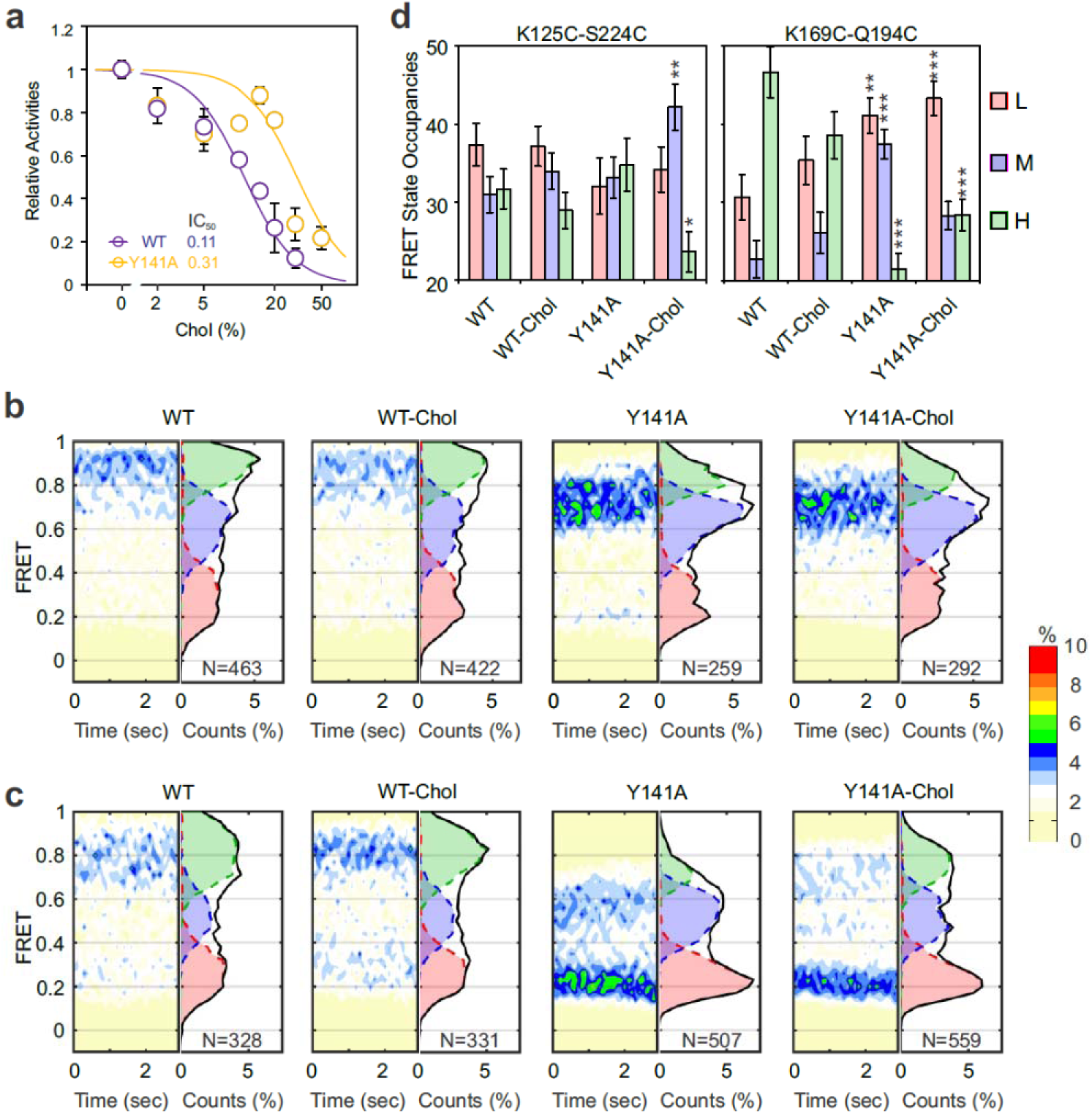
Y141A mutation enriches an intermediate S4 conformation to attenuate cholesterol inhibition. **a**. Dose-dependent inhibition of hHv1 WT and Y141A mutant fitted with the Hill equation. All data were presented as mean±SE, n=3-9. With the Hill coefficients fixed at 2, the half-inhibition concentrations (IC50) are 11% and 31% for WT and Y141A, respectively. **b** and **c**. FRET contour maps and histograms of FRET traces from the K125C/S224C (**b**) and K169C/Q194C (**c**) labeling sites, on the WT or Y141A backgrounds, without or with 20% cholesterol. The N number indicated the total number of FRET traces included in the analysis. **d**. FRET state occupancies of FRET traces from the K125C/S224C (a) and K169C/Q194C (b) labeling sites, on the WT or Y141A backgrounds, without or with 20% cholesterol. A kinetic model containing 3 FRET states, including low (L), medium (M) and high (H) FRET states, was used to idealize FRET traces and error bars indicate uncertainties in FRET state occupancy data obtained via bootstrapping. Two-tailed Student’s t-tests were performed to examine changes in FRET state occupancies in comparison to the control condition, with * denoting p<0.05, ** denoting p<0.01 and *** denoting p<0.001.

### Structural dynamics of the S4 segment underlie desmosterol and Y141A effects

Our smFRET measurements revealed that the S4 segment transits among low (L), medium (M), and high (H) FRET states. To further quantify how desmosterol and the Y141A mutation alter the conformational transitions of the S4 segment to regulate channel function, we calculated equilibrium constants (lnK) for transitions between L, M, and H FRET states from the smFRET data at both labeling sites (Fig. 5a). Since our smFRET measurements were performed under resting membrane potential of −85 mV, the cholesterol has very minimal impacts on the equllibrium constants of all 3 transition types, including LM, MH and LH (Fig 5a). Desmosterol, however, significantly enhances the M to H and L to H transitions at the K125C-S224C labeling site while suppressing them at the K169C-Q194C site, as expected from the opposite FRET directions reported by the two labeling sites for the same S4 movement (Fig 5a). Interestingly, the Y141A mutation altered transition kinetics similar to those of desmosterol, but only in the K169C-Q194C labeling sites. Based on our results, we proposed a kinetic model in which desmosterol and the Y141A mutation promote transitions from the closed (C) and intermediate (I) states to the open state (O) conformations to overcome the effects of cholesterol, which stabilizes the closed state conformations of the S4 segment in the hHv1 channel (Fig. 5b).

**Fig 5.**
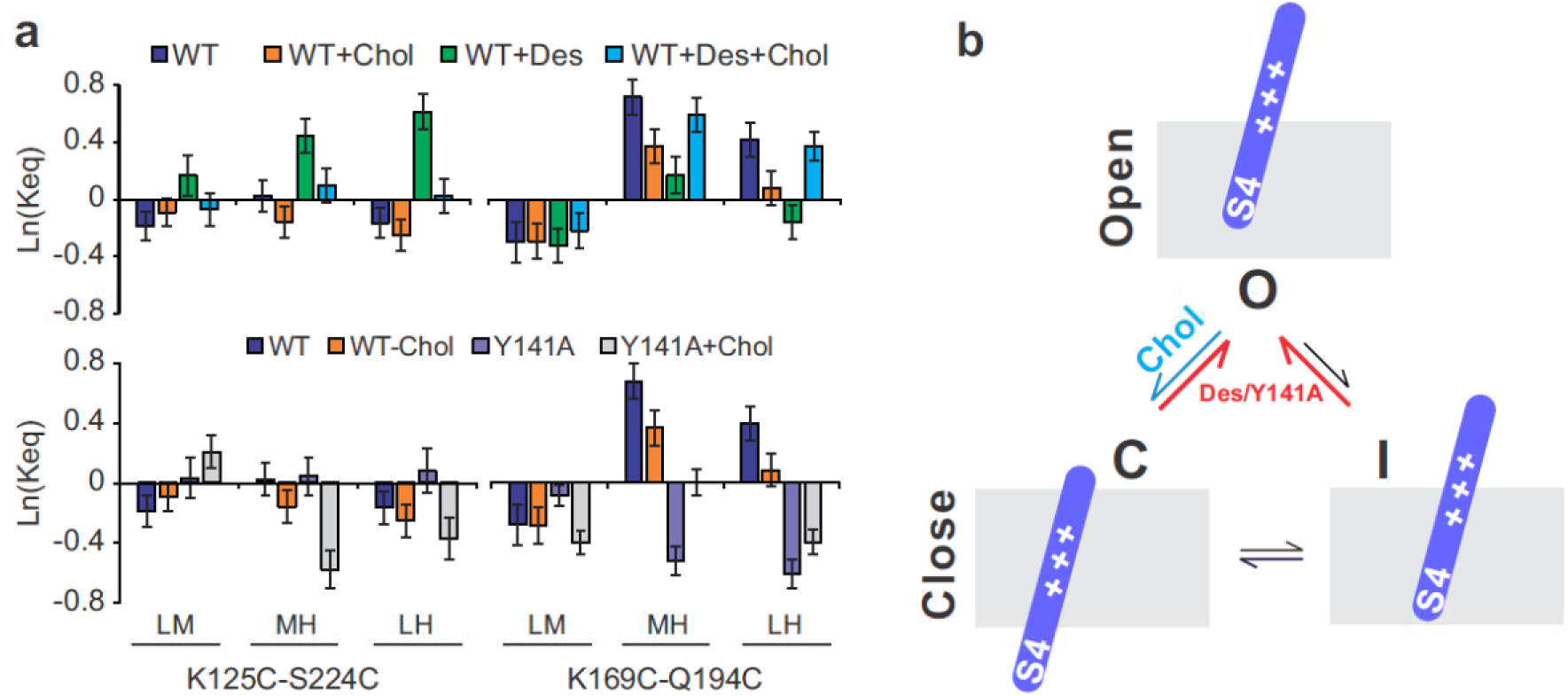
A kinetic model of the hHv1 channel regulated by membrane sterols. **a.** Effects of membrane sterols and the Y141A mutation on conformational transition kinetics of the hHv1 S4 segment. The equilibrium constants of low to medium (LM), medium to high (MH) and low to high (LH) FRET states were calculated from smFRET traces collected from the K125C-S224C and K169C-Q194C labeling sites, with error bars shown as propagated uncertainties of FRET state occupancy data. The smFRET data were collected at −85 mV to assess their effects on promoting the activated conformations of the S4 segment. **b**. A kinetic model of hHv1 channel gating regulated by membrane sterols. Cholesterol stabilizes the S4 segment at close state conformation (C), while desmosterol reverses cholesterol inhibition by promoting conformational transitions from close (C) to open (O) and intermediate (I) to open (O) states. As shown in panel **a**, desmosterol enhances the MH and LH transitions at the K125C-S224C labeling site and suppresses them at the K169C-Q194C labeling site. These opposite changes in transition kinetics are expected because the same S4 movement from close to open states is reported as increases in FRET at the K125C-S224C labeling sites but decreases at the K169C-Q194C labeling sites. The effects of the Y141A mutation are very similar to those of desmosterol.

## DISCUSSION

In our previous work, by reconstituting hHv1 proteins purified from *E. coli* host cells , we showed that cholesterol inhibits hHv1 channels, but the underlying mechanisms remained to be elucidated (15). In the present work, we further examined the effects of desmosterol, the immediate precursor to cholesterol, which, surprisingly, reverses cholesterol inhibition (Fig. 1d). The finding suggested that cholesterol metabolism may have a role in cellular pH homeostasis through regulating hHv1 channels, particularly under pathological conditions with unusual desmosterol levels (38, 39). It has been reported that overexpression of DHCR24, the enzyme that converts desmosterol to cholesterol, alleviates oxidative stress in many cell types by reducing ROS production (40–42). So, it would be very interesting to determine whether such effects result from regulating hHv1 channels by altering the cholesterol/desmosterol ratio in membranes. Desmosterol and cholesterol differ only in their side chains, with desmosterol carrying an additional double bond. The reversal of cholesterol inhibition by desmosterol suggested that the hHv1 channel can distinguish subtle structural differences between them. However, cholesterol and desmosterol also have distinct impacts on membrane biophysical properties, including lipid order, density, and flexibility (43, 44). Mutagenesis and docking simulation results suggested that cholesterol inhibits the hHv1 channel through multiple binding sites distributed across the transmembrane domain, rather than a single localized pocket. Our finding is consistent with emerging evidence that cholesterol can interact with ion channels at multiple, dispersed sites, as shown for inward-rectifier and BK potassium channels (23, 45, 46). Although the Y141A mutant exhibits activity similar to WT (Fig 4a), the mutation promotes a medium-FRET population in the S4 segment (Fig. 4b,c, d). As a result, even in the presence of 20% cholesterol, the conformational distributions of the S4 segment are not shifted significantly toward the resting state, represented by the low FRET population in the K125C-S224C and high FRET population in the K169C-Q194C labeling sites (Fig. 4b, c). These results align with our previous finding that cholesterol stabilizes the resting state conformations of the S4 segment (15) and further suggest that cholesterol inhibition of the hHv1 channel may be conformationally dependent. Based on the smFRET data, as shown in the kinetic model of Fig. 5b, cholesterol stabilizes the resting state conformations, while desmosterol promotes the open state conformations, and the Y141A mutation enriches a medium conformation, diminishing the effects of cholesterol (Fig. 5b). However, even with studies using stereoisomers of cholesterol and mutational studies, at this moment, we still can not rule out the possibility that cholesterol inhibition could result from both direct and indirect effects since studies have showed that the hHv1 channel and its major working partner, NADPH oxidases are both located in lipid raft domains rich in cholesterol (20, 22). So, it would be very interesting to study the localization of native hHv1 channels in primary cells and the effects of cholesterol depletion using super-resolution microscopy in the future.

## Acknowledgments

This work was supported by NIH grants 1R01GM142816 (SW), 1P50 MH122379 (DFC), and 2R35GM136409 (XZ), and by the University of Missouri-Kansas City (SW). The content of this article is solely the responsibility of the authors and does not necessarily represent the official views of the National Institutes of Health, the University of Missouri-Kansas City or the Washington University School of Medicine.

## Author Contributions

SW conceived the project and designed the studies; SH and SA performed the experimental research with help from Shuyi W and GW; RD and XZ performed computational modeling and simulation; MQ synthesized and DFC provided the ent-cholesterol and epi-cholesterol; SW and SH analyzed the experimental data; SW prepared the manuscript with input from all authors.

## Abbreviations

smFRET: single-molecule fluorescence resonance energy transfer
ATP: adenosine triphosphate
Cy3: Cyanine3
Cy5: Cyanine3
POPE: 1-palmitoyl-2-oleoyl-sn-glycero-3-phosphoethanolamine
POPG: 1-palmitoyl-2-oleoyl-sn-glycero-3-phospho-(1’-rac-glycerol)
ACMA: 9-Amino-6-Chloro-2-Methoxyacridine
CCCP: carbonyl cyanide m-chlorophenyl hydrazine
PEG: polyethylene glycol
NADPH: reduced nicotinamide adenine dinucleotide phosphate
DHCR24: 24-Dehydrocholesterol reductase

